# Context-dependent acetylation of the virulence regulator PhoP accounts for carbon-source specific intracellular growth program of *Mycobacterium tuberculosis*

**DOI:** 10.1101/2024.01.25.577197

**Authors:** Partha Paul, Khushboo Mehta, Rajat Ujjainiya, Bhanwar Bamniya, Bhuwaneshwar Thakur, Harsh Goar, Dibyendu Sarkar

## Abstract

*Mycobacterium tuberculosis* PhoP is essential for intracellular survival and virulence of the tubercle bacilli. Genetic evidences coupled with biochemical studies uncover that PhoP affects various aspects of *M. tuberculosis* pathophysiology including pH sensing during intracellular adaptation and carbon source utilization. Building on this observation, herein we report essentiality of the *phoP* locus in carbon-source specific mycobacterial growth. Further, our results on mycobacterial growth in the presence of different carbon sources suggest accumulation of acetyl CoA, a metabolic intermediate which acetylates major transcription factors. To explore the mechanism, we examined *in vivo* acetylation of PhoP, and our results suggest a link between acetylation of PhoP and mycobacterial carbon source utilization. Using a genetic screening, we identified PhoP-specific mycobacterial acetylases. Our two major findings that (a) acidic conditions of growth inhibit PhoP acetylation, which represses PhoP regulon by interfering with DNA binding activity of the regulator and (b) mycobacteria expressing acetylation-defective PhoP shows growth inhibition, together suggest a role of acetylation on mycobacterial growth via carbon-source utilization. These results have implications on intracellular survival and growth program of mycobacteria under varying environmental cues.

## INTRODUCTION

*M. tuberculosis* remains one of the most successful slow-growing human pathogens with a doubling time ranging from □ 20hours to 70 days either *in vitro*, in macrophages or in animal models [1]. During hypoxia, *M. tuberculosis* attains a state where the pathogen remains non-replicating, undergoing growth arrest but remain viable. It is believed that slow or arrested growth during nonreplicating persistence plays a key role in establishing a chronic infection and drug resistance. Both the macrophage phagosome and the caseum of granulomas indicate that *M. tuberculosis* is capable of integrating environmental signals like acidic pH and carbon source availability to fine-tune mycobacterial growth and metabolism. A previous report, along the line, elegantly showed enhanced growth of triacylglycerol (TAG) synthase mutant (*tgsI*) relative to WT bacilli under acidic pH [2]. More recent work by Baker et al provided clear evidence of slow growth under acidic pH and its control by host-associated carbon sources [3]. However, the underlying mechanism is yet to be clearly understood.

The hallmark feature of *M. tuberculosis* metabolic plasticity includes the ability to co-catabolize multiple carbon sources simultaneously [4]. In fact, reprogramming of carbon metabolism is believed to play a key role in mycobacterial adaptation inside the host, and consequently, a number of studies have identified major mechanisms of mycobacterial carbon metabolism during infection [4–8]. In an intracellular milieu using the flexible metabolic network, a minor fraction of *M. tuberculosis* moves to a reduced growth-state to adapt to a range of environmental cues including stresses, like nutrient starvation, hypoxia or antibiotic treatment[9–16] . Consequently, in a non-replicating state *M. tuberculosis* remains less susceptible to the antimicrobial impacts of the environmental stress conditions [17, 18]. This adaptive strategy primarily relies on the ability of the bacilli to switch between the replicating and nonreplicating state via rerouting of its metabolic fluxes.

Several studies have documented that *M. tuberculosis* growth and regulated gene expression are coupled by pH in the surrounding environment [1, 2, 19–21]. In keeping with this, a) *M. tuberculosis* is known to encounter acidic environment *in vivo*, b) bacterial adaptative response under acidic pH remains critically important for pathogenesis, and c) transcription of *M. tuberculosis* genes in response to acidic conditions of growth, both *in vitro* and in macrophages, reveals significant variation of global gene expression [1, 20, 22]. Strikingly, many acid-inducible mycobacterial genes are also closely connected with carbon metabolism. For instance, *pks2*, *pks3*, and *pks4* genes are linked to the biosynthesis of complex cell envelope lipids like sulpholipids, diacyl, and polyacyltrehalose (SLs, DATs and PATs) [23–25]. Likewise, expression of *icl1* (isocitrate lyase), which is required for infection, suggesting that *M. tuberculosis* metabolizes fatty-acid derived Acetyl CoA [5] or cholesterol-derived propionyl CoA [9], shows significant induction under low pH. Furthermore, the acid and phagosome-regulated locus *aprABC*, which is significantly induced under acidic pH [2], remains linked with the regulation of triacylglycerol (TAG) accumulation and governs carbon and propionate metabolism genes [26]. Remarkably, all of these genes are regulated by the PhoP-PhoR regulatory system, require PhoP for its expression, and in most cases the corresponding promoters are strongly bound by the response regulator [23, 27, 28]. However, the mechanism by which PhoP connects between the two environmental cues, acidic pH and carbon-source availability, remains unknown.

The objective of this study was to identify the underlying mechanism of carbon source-specific metabolic shift, that promotes growth arrest or persistence under acidic environment. Herein, we show here that the critical two-component regulator PhoP, which plays a key role in modulating pH driven adaptation to the macrophage phagosome [20, 29], functions to connect the two environmental cues within the host, acidic pH and carbon source availability. In this connection, we demonstrate that PhoP undergoes enzymatic acetylation at a conserved lysine residue, and represses *phoP* regulon. Although acetylation of PhoP is inhibited under acidic conditions of growth in rich medium, acetate as a model fatty acid in the growth medium promotes acetylation of the regulator via a pH-independent mechanism. Collectively, these results uncover that in a carbon source-dependent manner PhoP undergoes acetylation, represses its own regulon, and triggers a global metabolic rearrangement which promotes mycobacterial growth arrest and/or persistence.

## Results

### Mycobacterial growth in the presence of acetate as a carbon source under low pH conditions depend on the *phoP* locus

Although *M. tuberculosis* displays growth arrest under low pH in a carbon-source specific manner [3, 30], the molecular basis of growth arrest/inhibition remains poorly understood. To examine the impact of available carbon-source on mycobacterial growth under low pH and normal conditions, we grew WT-H37Rv *in vitro* using glycerol and acetate, as specific carbon sources, respectively, both under normal and acidic conditions of growth (Fig. 1A). Upon enumeration of CFU values, we observed that WT bacilli shows a significantly reduced growth under pH 5.5 relative to pH 7.0, when glycerol was used as the carbon source. In contrast, WT-bacilli displayed a comparable growth at pH 5.5 and pH 7.0, when acetate was used as the carbon source. These results are consistent with previous results suggesting mycobacterial failure to utilize glycerol as a carbon source under acidic pH [3]. Additionally, these results clearly demonstrate that acetate can be used as a mycobacterial carbon source with comparable efficiency, both at the normal and acidic pH (Fig. 1A). Previous studies uncovered that mycobacterial *phoP* locus plays a major role in intra-phagosomal adaptive response under acidic pH [20, 29]and impacts on carbon source utilization [3]. In fact, a transposon mutant of *M. tuberculosis* strain carrying mutations in *phoP* was shown to be effectively growing in pyruvate under acidic conditions of growth relative to WT bacilli [3]. Thus, we compared *in vitro* growth of WT-H37Rv and a *phoP*-KO mutant (Δ*phoP*-H37Rv) in the presence of glycerol and acetate (as carbon source) both under normal and acidic pH, respectively. Our results demonstrate that both WT-H37Rv and *phoP*-KO mutant were unable to utilize glycerol under acidic pH (Fig. 1B). However, under normal conditions (pH 7.0), WT bacilli showed better *in vitro* growth relative to the *phoP*-KO mutant (Fig. S1A). In contrast to bacterial growth under acidic pH in the presence of glycerol as the carbon source, WT bacilli showed a significantly higher growth compared to *phoP*-KO mutant in the presence of acetate and under acidic conditions of growth (Fig. 1C). It should be noted that a comparably similar and significant growth defect of *phoP*-KO mutant was not apparent under normal conditions (pH 7.0) of growth in acetate (see Fig. S1B). These results suggest importance of the *phoP* locus in utilizing acetate as the carbon source under acidic conditions of growth. In keeping with the above results, additional growth curve experiments further confirm that while mycobacteria fail to effectively utilize glycerol as the carbon source under acidic conditions of growth (Fig. 1D), WT-H37Rv but not *phoP*-KO mutant, can effectively utilize acetate as the carbon source under acidic conditions of growth (Fig. 1E).

**Fig. 1:**
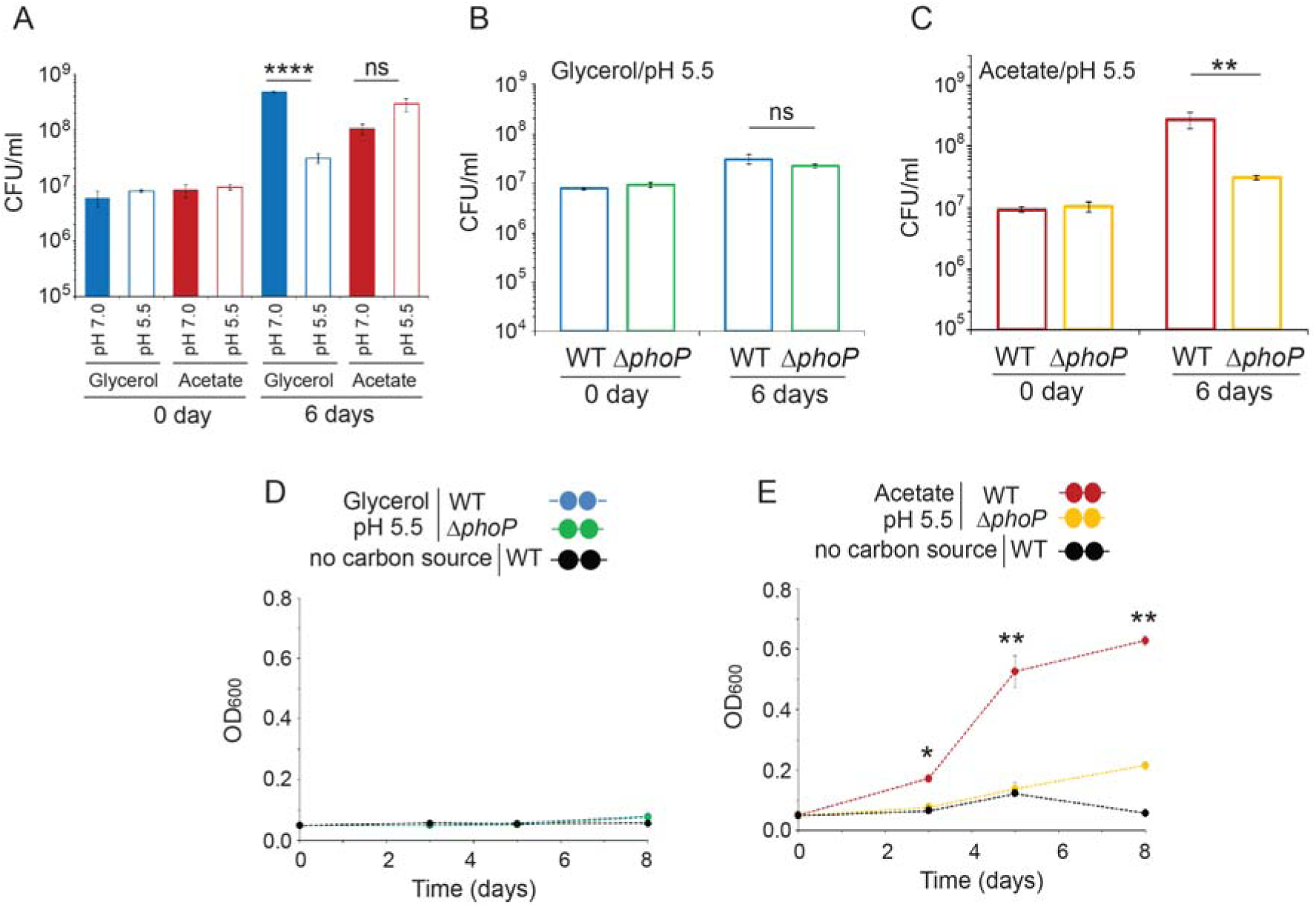
*phoP* is required for utilization of acetate as a carbon source. *Mycobacterium tuberculosis* was grown in a minimal media containing either glycerol or acetate, under acidic conditions (pH 5.5). In this experiment, indicated mycobacterial strains were inoculated at an initial OD_600_ of 0.05 at specific media, as described in the Methods, and growth was monitored at different time points to generate the growth curve. To enumerate the CFU values, cells were harvested both at the onset of growth (0 day) and on the 6^th^ day. The experiments represent average of two biological repeats with at least one technical repeat (**P□≤□0.01).

### M. tuberculosis PhoP is acetylated in vivo

Notably, acetate upon its utilization as a carbon source is efficiently converted to acetyl-CoA, which in conjunction with acetyl transferase often acetylates major transcription factors. With growing evidence suggesting a major role of the *phoP* locus in various mycobacterial stress response including acidic pH, we investigated *in vivo* acetylation of PhoP for cells grown in 7H9 medium with carefully controlled varying single stress conditions (see methods for details) (Fig. 2A). In this assay, His-tagged PhoP was expressed from p19kpro [31] in WT-H37Rv and acetylation of PhoP was probed using anti-acetyl antibody. Also, cell extracts from identical samples were probed with anti-His antibody to confirm presence of PhoP at comparable amounts. PhoP was found to be acetylated under normal as well as varying stress conditions like oxidative stress or salt stress. However, we observed a significant decline in PhoP acetylation for mycobacterial cells grown under acidic conditions. These results suggest that independent of carbon source, acidic condition of growth inhibits *in vivo* acetylation of PhoP. We next studied *in vivo* acetylation of PhoP using glycerol and acetate as carbon sources in the growth media (Fig. 2B). Our results clearly demonstrate that in the presence of glycerol as carbon source, PhoP is insignificantly acetylated even under normal (pH 7.0) conditions of growth (lane 1, Fig. 2B). Note that in the absence of detectable *in vitro* growth we were unable to determine level of acetylation of PhoP for cells grown in presence of glycerol under acidic pH. In contrast, in the presence of acetate as the carbon source, PhoP was comparably acetylated regardless of growth conditions (compare lanes 2 and 3, Fig. 2B). These results suggest that PhoP acetylation is linked to mycobacterial growth under acidic conditions in presence of acetate as the carbon source.

**Fig 2:**
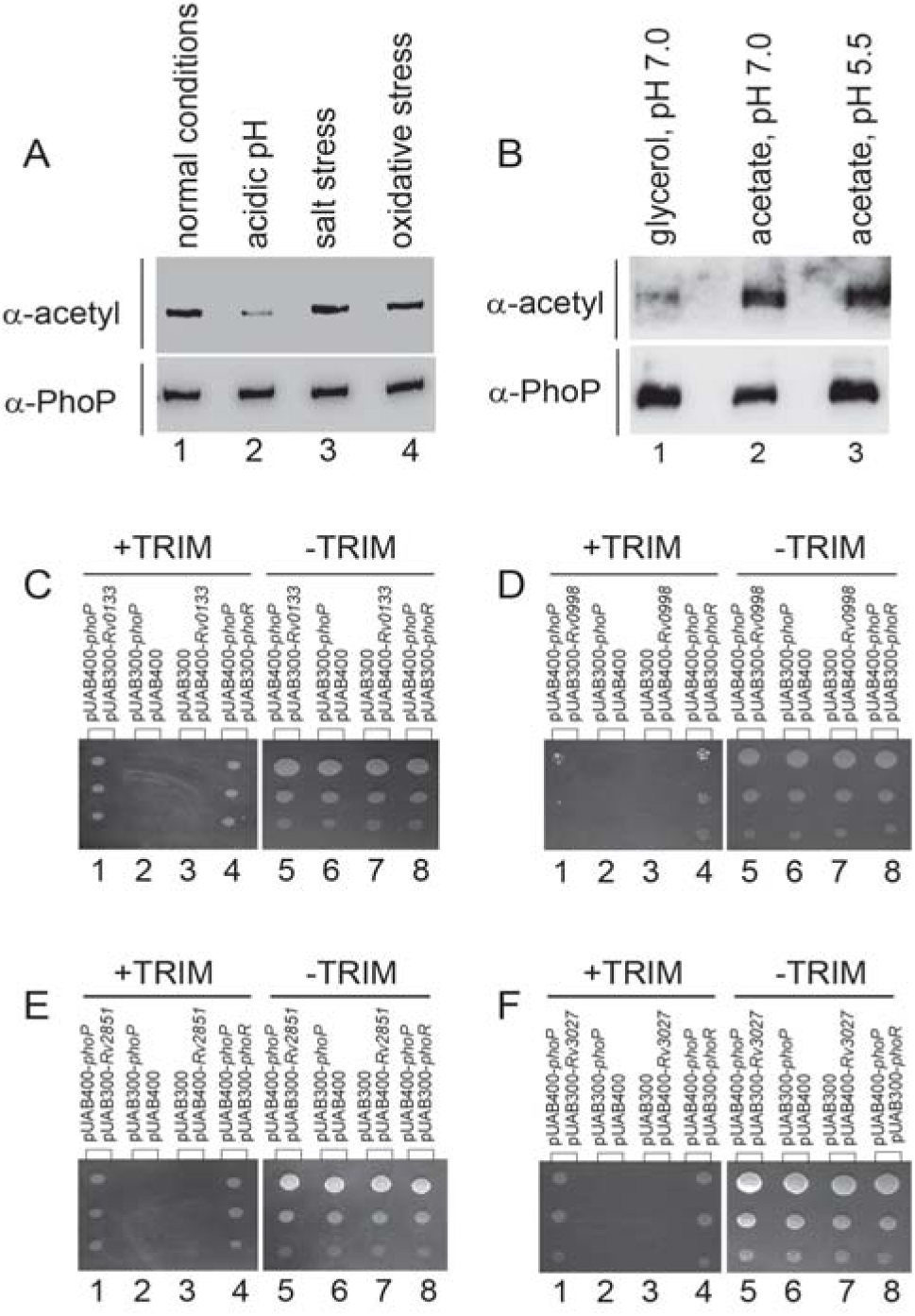
*In vivo* acetylation of PhoP under different stress conditions. (A-B) *M. tuberculosis* ectopically expressing His-tagged PhoP (see methods for details) was grown under defined stress conditions. Next, PhoP was purified from each of the crude lysates using Ni-NTA affinity chromatography, and immunoblotted to assess acetylation using anti-acetyl antibody. As a control, expression of PhoP was compared in identical samples using anti-PhoP antibody. The blots are representative of at least two independent experiments. (C-F) To probe interaction of PhoP with mycobacterial acetyltransferases, we utilized mycobacterial protein fragment complementation (M-PFC) -based genetic screen (see methods for details). Importantly. *M. smegmatis* expressing *M. tuberculosis* protein pairs (C) *phoP/Rv0133*, (D) *phoP/Rv0998,* (E) *phoP/Rv2851, or* (F) *phoP/Rv3027* displayed growth in presence of TRIM. However, empty vector controls under identical conditions did not grow in the presence of TRIM, suggesting interaction between the specific acetyltransferases and PhoP. Note that *phoP/ phoR* pair was used as a positive control, and plates lacking TRIM showed comparable growth of all the strains used in this assay.

Having shown that PhoP is acetylated *in vivo*, using a bacterial two hybrid system we next probed for the specific acetyl transferase(s) which likely acetylate the master regulator. In this experiment, we utilized a protein fragment complementation (M-PFC) assay using *M. smegmatis* [32]. The objective was to detect interactions between PhoP and major mycobacterial acetyl transferases (Figs. 2C-F), which acetylate mycobacterial proteins. According to this assay, two mycobacterial proteins which likely interact to each other, are fused to C-terminal domains of complementary fragments of mDHFR. Subsequently, the bacterial cells harboring both proteins display bacterial resistance to trimethoprim (TRIM) as a result of functional reconstitution of active and full-length DHFR protein. Thus, *M. smegmatis* strains harbouring the appropriate constructs were grown on 7H10/Kan/Hyg in the presence or absence of 10 µg/ml TRIM. Strikingly, *M. smegmatis* cells co-expressing 4 out of 11 acetyl transferases (Rv0133, Rv0998, Rv2851, Rv3027) and PhoP protein exhibited growth in TRIM containing plates, whereas none of the other acetyl transferases under identical conditions, showed *in vivo* interaction with PhoP (Fig. S2, panels A-G). As controls, all of the co-transformants harbouring acetyl transferase/PhoP pairs showed comparable bacterial growth in plates lacking TRIM. From these results, we conclude that PhoP acetylation is attributable to specific mycobacterial acetyl transferase(s).

### *M. tuberculosis* PhoP is a substrate for Rv0998, Rv2851, and Rv3027 acetyl transferases

Having shown that *M. tuberculosis* PhoP undergoes *in vivo* acetylation and specific acetylases interact with PhoP, we wished to examine *in vitro* acetylation of the regulator by the interacting acetylases. To this objective, four PhoP interacting acetyl transferases were cloned, expressed and purified from *E. coli* and the recombinant acetyl transferases were incubated with purified PhoP in acetylation mix containing acetyl CoA. To probe acetylation, samples were resolved by SDS-PAGE and immunoblotted with anti-acetyl antibody (Fig. 3). Identical samples were also blotted with anti-His antibody to detect PhoP and corresponding acetyl transferases used in the assay. Importantly, we observed insignificant acetylation of PhoP in the absence of any acetyl transferase protein (lane 1, Fig. 3A). Similarly, in the presence of acetyl-CoA alone we were unable to find significant acetylation of the regulator (lanes 2-3, Fig. 3A). However, lanes 4-7 showed detectable PhoP acetylation in the presence of Rv0133, Rv0998, Rv2851 and Rv3027 acetyl transferases, respectively. More robust acetylation was detectable in lanes 5-7 containing Rv0998, Rv2851, and Rv3027, respectively. As a control, acetylation of recombinant DosR was investigated using an identical set up (Fig. 3B). Under identical conditions examined, Rv0133 showed robust acetylation of recombinant DosR (lane 4, Fig. 3B), whereas other acetyl transferases under identical conditions were largely ineffective. These, these results suggest that PhoP is a specific substrate for acetylation by Rv0998, Rv2851, and Rv3027 acetyl-transferases.

**Fig. 3:**
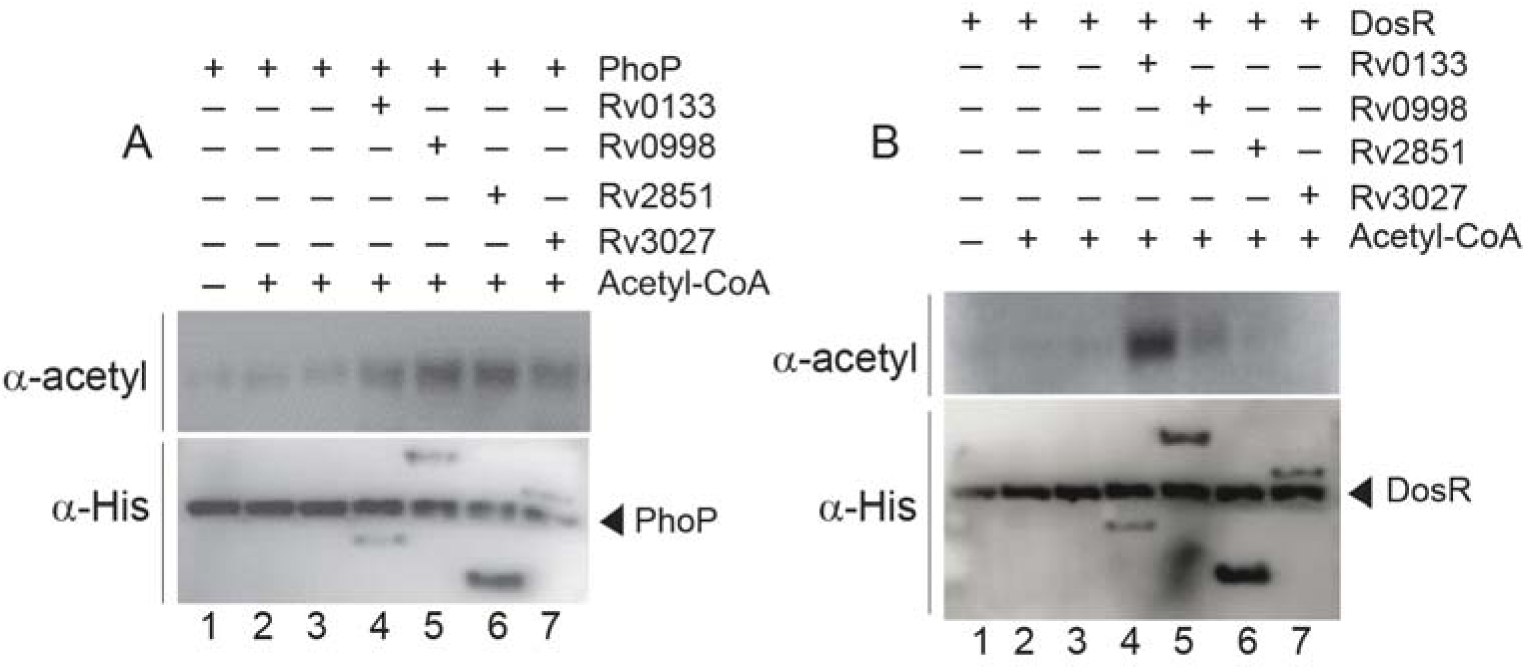
*In vitro* acetylation of PhoP by acetyltransferases. In this assay, ≈ 2 µg of recombinant PhoP (A) or DosR (B) was incubated with or without purified acetyltransferases (≈ 1 µg) in acetylation mix containing 0.5 mM acetyl-CoA, and incubated at 25°C for 2 hours. The reaction products were analyzed by SDS-PAGE, and probed through Western blot analysis using anti-acetyl antibody. Presence of comparable amounts of PhoP was ensured by Western blot analysis of identical samples using anti-His antibody. The blots are representative of at least two independent experiments.

### Acetylation of PhoP represses *phoP* regulon

*M. tuberculosis* PhoP, as a major transcription factor, regulates expression of □2% of the genome. To investigate whether acetylation impacts activity of the transcription factor, expression of the *phoP* regulon genes was examined by real-time qPCR using mycobacterial cells grown in glycerol and acetate as carbon sources, respectively. The results show that PhoP-regulated genes are significantly down-regulated in mycobacterial cells grown in acetate compared to mycobacterial cells grown in glycerol (Fig. 4A). It should be noted that expression level of PhoP remains largely comparable in these cases (see Fig. 2B). Thus, these results suggest that it is the PhoP-activity, and not the level of intra-mycobacterial level of PhoP, which affects expression of the regulon. Given the fact that PhoP is hyper-acetylated (>10-fold, based on limit of detection in these assays) when mycobacterial cells are grown in acetate relative to glycerol, these results strongly suggest that PhoP acetylation inhibits transcription of PhoP regulon genes.

**Fig. 4:**
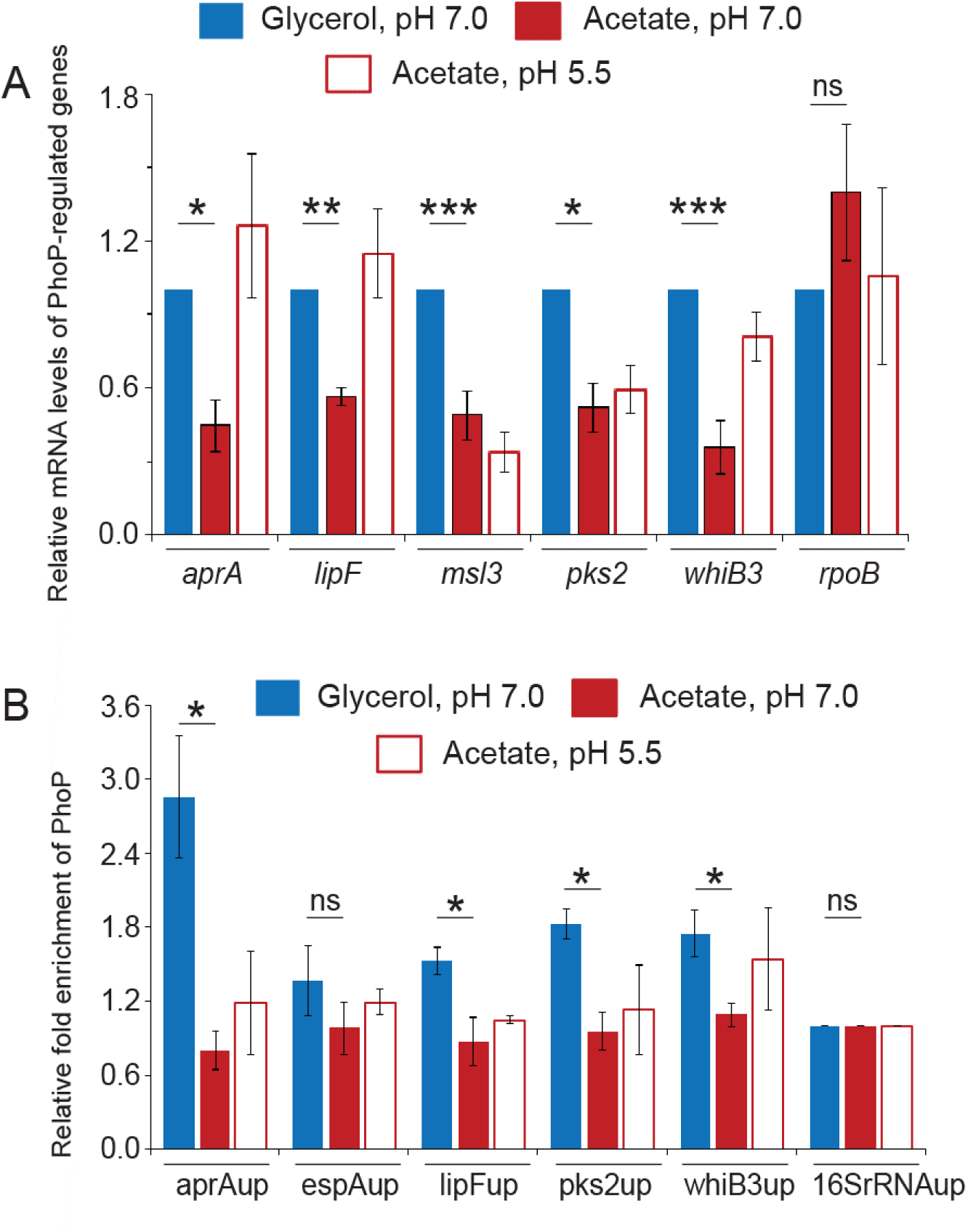
Mycobacterial growth conditions with varying carbon sources impact expression of the PhoP regulon. (A) WT-H37Rv was either grown in glycerol and acetate under normal pH or in acetate under low pH (acidic conditions) till log phase. Next, RT-qPCR assays using total RNA isolated from mycobacterial cells and appropriate primers examined expression levels of the *phoP* regulon genes. Note that 16s rRNA was used as an endogenous control, and the average fold difference represents technical repeats with at least two independent RNA preparations. (B) For ChIP assays, the cells lysates were immunoprecipitated using anti-PhoP antibody, and qPCR using appropriate (upstream region-specific) primers determined the fold enrichments relative to the mock signal (no antibody control). 16SrDNAup was used as the endogenous control. The experiments were performed with technical repeats using at least two biological samples (*P < 0.05; **P < 0.01; ***P < 0.001).

To verify whether DNA binding activity of PhoP is impacted, we next performed chromatin immunoprecipitation assays using anti-PhoP antibody, followed by qPCR (Fig. 4B). Our ChIP-qPCR results demonstrate that for mycobacterial cells grown in acetate, PhoP is recruited within target promoters to a significantly lower extent compared to cells grown in glycerol. Coupled with the fact that PhoP expression remains comparable regardless of the carbon source used (Fig. 2B), these results suggest that specific acetylation inhibits effective DNA binding activity of the regulator. As acetylation of PhoP is promoted in the presence of acetate as the carbon source (relative to glycerol) with synthesis of acetyl-CoA as a metabolic intermediate, these results suggest that inhibition of DNA binding activity of PhoP is attributable to PhoP acetylation.

### Acetylation of PhoP at K195 regulates carbon-source specific mycobacterial growth

To investigate which lysine residue of PhoP undergoes acetylation, bands containing purified PhoP protein was excised out from acrylamide gel and subjected to mass spectrometry analyses. Among many other lysine residues, K195 was the most frequently acetylated site found in replicate samples with ∼99% confidence (Fig. 5A). Critically, a 41-Da increase in the mass of the peptide (^1^YFVINAGTVLSK^12^PK) was consistent with acetylation of the K12 residue of the peptide, suggesting K195 as the primary acetylation site of the regulator. The LC/MS/MS spectra of the peptide with modified fragment ions (K195Ac) is shown in Fig 5A. This result is consistent with previously reported global acetylome data of *M. tuberculosis*, suggesting K195 of PhoP as the primary site of acetylation [33]. Notably, amino acid sequence alignment of the PhoP family of regulator uncovers K195 as one of the four conserved lysines of PhoP (Fig. S3A).

**Fig. 5:**
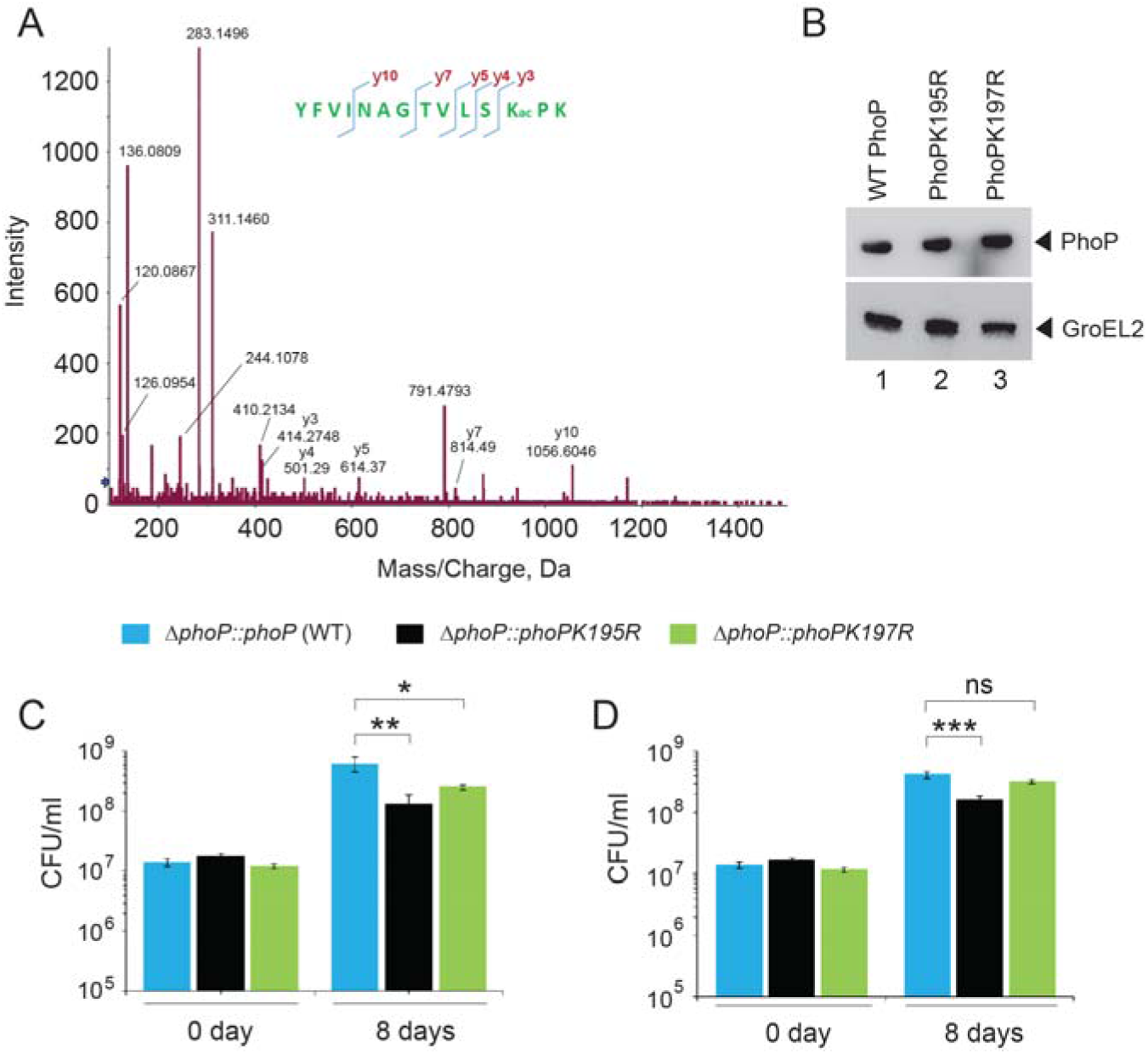
*M. tuberculosis* PhoP is acetylated at Lys195. (A) Mass spectrometry analysis of PhoP-derived tryptic peptides. 6xHis-tagged PhoP was analyzed with LC/MS/MS after trypsin digestion. Shown is the spectrum covering the region from 100 to 1500 m/z including the peptide comprising the acetylated K195. Acetylation was observed at K12 residue of the peptide ^1^YFVINAGTVLSK^12^PK. The ESI-MS/MS spectra obtained for the peptide with CID of +2 m/z ion at 789.9429 is shown in the figure. The modified y-ions showing an increase of ∼41-Da in the peptide mass is annotated and the plot is representative of multiple independent experiments. (B) Wild-type and indicated mutants of PhoP were expressed in the *phoP*-KO mutant *M. tuberculosis* as described in the ‘methods’ section, and expression of PhoP was probed by immunoblotting. GroEl2 was used as a loading control. (C-D) Utilization of acetate as the carbon source by mycobacteria under normal (C) and acidic conditions (D) of growth is determined by the acetylation of PhoP at K195. Mycobacteria harboring wild-type or indicated *phoP* mutants were inoculated at an initial OD_600_ of 0.05 in the presence of acetate as the carbon source. To enumerate the CFU values, cells were harvested at the onset of growth (0 day) and on the 6^th^ day. The experiments represent average of two biological repeats and at least one technical repeat (*P<0.05; **P<0.01; ***P<0.001).

To investigate the impact of PhoP acetylation on mycobacterial growth, we next constructed an acetylation-defective PhoP mutant (PhoPK195R). As a control, we also included PhoPK197R replacing K197 of PhoP with Arg (see methods for details). These mutant proteins were expressed in the *phoP*-KO background, and Fig. 5B shows that under the conditions examined, both the WT and the mutant proteins are expressed at a comparable level. Next, during *in vitro* growth experiments with acetate as the carbon source, we observed a significantly lower growth of mycobacterial strain expressing acetylation-defective PhoP relative to acetylation-proficient mutant or WT bacilli both under normal and acidic conditions of growth (Figs. S3B and S3C). Consistent results were obtained upon CFU enumeration (Figs. 5C and 5D). However, a significantly less pronounced impact on growth was observed for mycobacterial strain expressing PhoPK197R, which can undergo acetylation. These results suggest that acetylation of PhoP is linked with mycobacterial growth arrest under acidic pH in the presence of acetate as the carbon source.

Having shown that K195 residue, the primary acetylation site of PhoP plays a major role during *in vitro* growth of the tubercle bacilli under acidic pH, we used WT and the mutant strains to infect murine macrophages (Fig. 6A). Although mycobacteria expressing wild-type PhoP could effectively inhibit phagosome-lysosome fusion, remarkably the strain expressing PhoPK195R readily matured into phagolysosomes, strongly indicating increased trafficking of the mycobacterial strain to lysosomes. As a control, mycobacteria expressing PhoPK197R inhibited phagosome-lysosome fusion with comparable efficiency as that of WT bacilli. Together these results including bacterial co-localization data (Fig. 6B) and Pearson’s plot (Fig. 6C) suggest that the K195 residue of PhoP plays a major role during intracellular growth of mycobacteria. In keeping with these results, 48 hr post infection mycobacteria expressing mutant PhoPK195R showed a significantly reduced intra-phagosomal growth relative to bacterial strains expressing either wild-type PhoP or the mutant PhoPK197R (Fig. 6D), suggesting that K195 residue of PhoP remains essential for mycobacterial growth in macrophages.

**Fig. 6:**
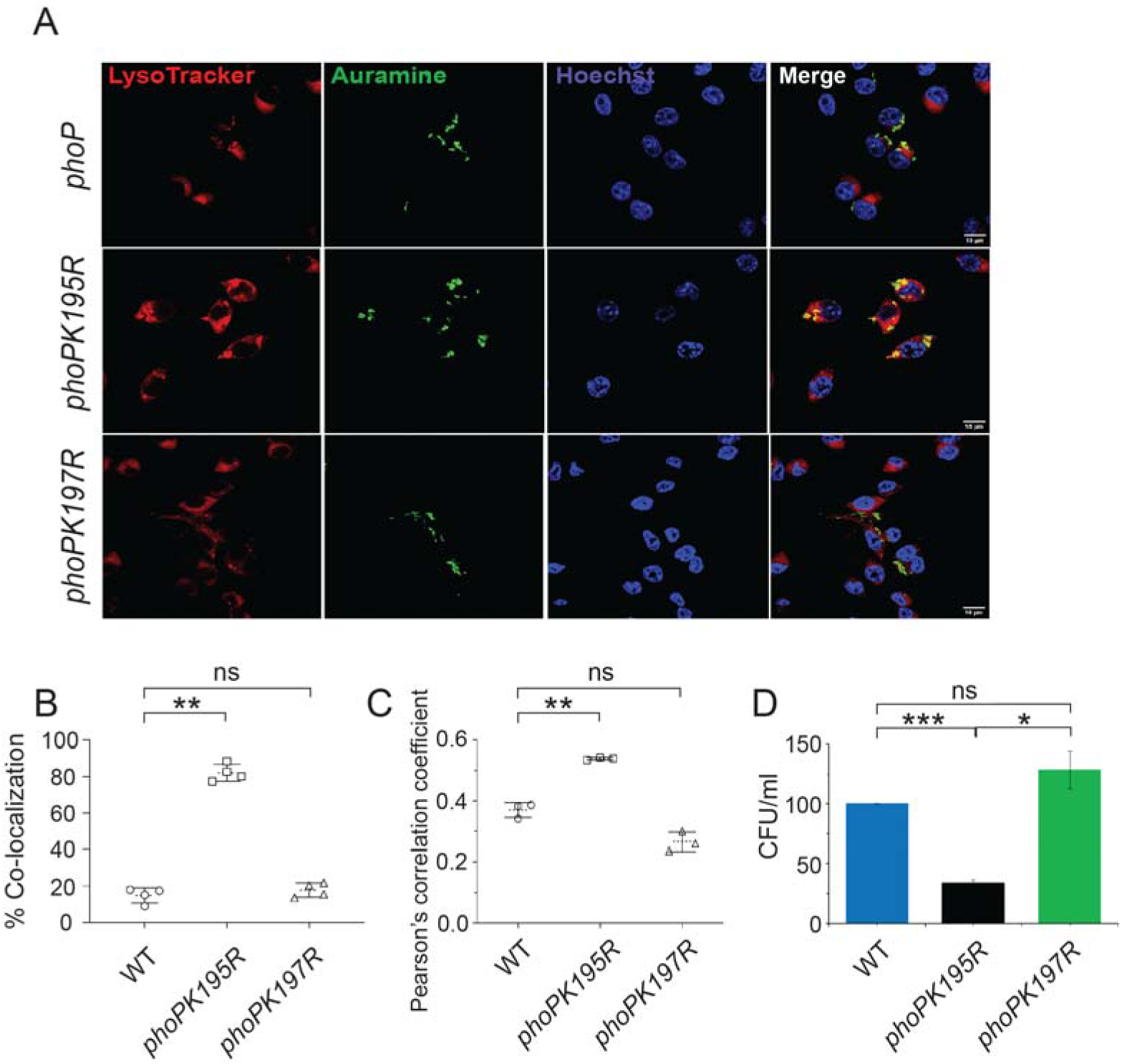
K195 residue of PhoP contributes to mycobacterial survival in cellular models. (A) Indicated *M. tuberculosis* strains expressing wild-type and mutant PhoP proteins were used to infect murine macrophages. Mycobacteria and host cells were stained with phenolic auramine solution, and LysoTracker respectively. Host cell nuclei were made visible by Hoechst dye. Three fluorescence signals (Mycobacterial strains: green; lysosomes: red and host nuclei: blue) and their merging are displayed by confocal images (scale bar: 10 µm). (B) Co-localization of auramine labelled mycobacterial strains with Lysotracker was investigated by visually scoring yellow and green punctas from at least 50 infected cells originating from 10 different fields of each of the 4 independent experiments. To determine percent co-localization, the number of yellow punctas were divided by the total number of punctas (yellow plus green) as described previously [49]. The results display average values from 50 infected cells (n=50) of each independent experiment with standard deviations from four biological replicates (**P≤ 0. 01). (C) The data present Pearson’s correlation coefficient of images displaying internalized auramine-labelled mycobacteria and Lysotracker red marker in macrophages, and were evaluated using image-processing software NIS elements (Nikon). Average values with standard deviations were obtained from three independent experiments (*P<0.05; **P<0.01). (D) To examine contribution of residue K195 to mycobacterial survival in cellular models, murine macrophages were infected with indicated mycobacterial strains, and 72-hour post infection intracellular bacterial CFU (normalized to 3-hour post infection) were enumerated. The results show average values from biological duplicates (*P<0.05; ***P<0.001).

## Discussion

Our present understanding on host-related *M. tuberculosis* metabolism is mostly organized around central carbon metabolism [34], which represents a key determinant enabling the bacilli to replicate and persist within the host. Thus, understanding the metabolic pathways which are utilized by mycobacteria to adapt to host-guided carbon source availability remains critically important to TB pathogenesis. In this connection glyoxylate shunt has drawn attention as *M. tuberculosis* utilizes this pathway to support its *in vivo* growth via fatty acid metabolism [5]. In this study, we investigated the role of carbon metabolism in mycobacterial growth control or persistence. We used acetate as a model fatty acid since host-derived fatty acids remain the most relevant carbon source for intracellular mycobacteria during acute and chronic phases of infection [5, 11, 35].

Given the link that exists between *M. tuberculosis* PhoP and its pH-driven adaptation to the macrophage phagosome in one hand [21, 29], and between variable carbon source utilization and requirement of the *phoP* locus on the other hand [3, 21, 26], we sought to identify the mechanism which connects the two environmental cues, namely acidic pH and mycobacterial carbon-source utilization. Herein, we show that mycobacterial utilization of acetate as a carbon source requires the presence of the *phoP* locus. Further, we demonstrate that PhoP undergoes enzymatic acetylation at a conserved lysine residue, inhibits DNA binding functions of the transcription regulator and represses the *phoP* regulon. Although acetylation of PhoP is inhibited under acidic conditions of growth in rich medium, acetate in the growth medium promotes acetylation of the regulator via a pH-independent mechanism. In keeping with these findings, mycobacterial strain expressing an unacetylated PhoP homolog remains incapable to utilize acetate under acidic conditions of growth and exhibits growth defect both *in vitro* and in macrophages. Collectively, these results uncover that in a carbon source-dependent manner PhoP undergoes acetylation, represses its own regulon, and triggers a global metabolic rearrangement which promotes mycobacterial persistence.

Although phosphorylation of PhoP plays an important role to activate or repress gene expression *in vivo* [23], regulation of functioning of the transcription factor by N^€^-lysine acetylation remains largely unknown. To circumvent the problem arising due to influence of other factors impacting mycobacterial growth, we first identified *phoP*-dependent mycobacterial growth in the presence of acetate, a model fatty acid as the carbon source. It should be noted that fatty acids, which remain the most relevant carbon source in an intracellular milieu [35, 36] often leads to generation of acetyl CoA as the metabolic intermediate. Thus, we probed for *in vivo* acetylation of PhoP in mycobacterial cells grown in the presence acetate as the carbon source under normal and acidic conditions of growth. In keeping with acetylation data, we next identified PhoP-specific mycobacterial acetyl transferases which acetylates PhoP in the presence of acetyl-CoA (Fig. 3), and PhoP in the acetylated form represses *phoP* regulon (Fig. 4). These results are consistent with identifying the conserved and homologous lysine residue (K195) of PhoP as the primary site of acetylation. Considering the location of the positively charged residue proximal to the DNA binding helix-turn-helix motif of PhoP [37, 38], we speculated that acetylation of K195 may regulate DNA binding activity of the regulator. To examine this possibility, K195 was mutated to arginine (R) since the conservative substitution maintains the positive charge, and at the same time represents an unacetylated homologue of the regulator. In conclusion, we observed a striking similarity in growth inhibition between *phoP*-KO mutant and a mycobacterial strain expressing unacetylated homolog of PhoP, both *in vitro* and *in vivo*. These results coupled with our studies on acetylation of PhoP suggest that among the reasons for mycobacterial growth arrest is a single conserved K195 residue of the virulence regulator which undergoes N^€^-lysine acetylation. What offers a new mechanistic insight is the finding that acetylation of Lys-195 of PhoP is critically important for carbon-source specific growth, which in turn, regulates mycobacterial persistence. This is shown by our growth experiments using mycobacterial strains expressing acetylation-proficient and unacetylated homolog of PhoP (Figs. 5 and 6). Thus, PhoP contributes to regulation of carbon-source dependent mycobacterial growth under acidic pH, facilitates an integrated view of our results. However, these results do not explain *phoP*-independent mycobacterial growth inhibition in the presence of glycerol and under acidic pH. Nonetheless, our findings establish a novel regulatory link between PhoP acetylation with host-related carbon source utilization impacting mycobacterial persistence.

It is noteworthy that WT-bacilli displays higher drug tolerance when cultured in acetate relative to glycerol medium, suggesting that metabolic network required for acetate metabolism is linked to mycobacterial drug tolerance [39]. The findings that acidic conditions of growth in one hand inhibits PhoP acetylation leading to repression of *phoP* regulon, and on the other hand stimulates PhoP phosphorylation which activates PhoP regulon, tempt us to speculate that strikingly contrasting outcomes by the two different post-translational modifications of PhoP presumably maintain the fine balance of activation status of *phoP* regulon, which indeed is a global regulator of metabolic network contributing to comprehensive stress response and pathogenicity of the tubercle bacilli.

## Materials and Methods

### Bacterial strains, media, and growth conditions

Cloning of various ORFs and expression of recombinant proteins utilized *E. coli* DH5α and *E. coli* BL21 (DE3), respectively. *E. coli* strains were grown at 37°C with shaking and transformants were selected in presence of appropriate antibiotics. For growing mycobacterial cultures, *M. tuberculosis* H37Rv and *M. smegmatis* mc^2^155 were grown at 37°C in Middlebrook 7H9 broth (Difco) media or 7H10 agar. *M. tuberculosis* lacking *phoPR* operon (*ΔphoP*-H37Rv) and the complemented strain have been mentioned previously [24]. *M. tuberculosis* strains were made competent, transformed and selected in antibiotics containing plates [40]. To grow mycobacteria in media containing single carbon source, minimal media (pH 7.0/5.5) was used as described [3, 30], and either glycerol or acetate at a concentration of 10 mM was supplemented as the nutrient carbon source. Cultures at the onset (0 day) and after certain time (6 day) were serially diluted, and plated at specific dilutions to enumerate CFU. For *in vitro* growth under a specific stress, indicated mycobacterial strains were grown to mid-log phase (OD_600_ 0.4 to 0.6) and exposed to different stress conditions as described previously [41].

### Cloning

Recombinant PhoP and DosR were cloned, and expressed as His_6_-tagged proteins in *E. coli* as described previously [42, 43]. *M. tuberculosis* acetyltransferase encoding ORFs *rv0133* and *rv0998* were cloned between PstI/HindIII, and NcoI/HindIII sites of pET-28b, respectively. Similarly, ORFs encoding *rv2851* and *rv3027* were cloned between BamHI/HindIII sites of pET-28a. His_6_-tagged PhoP was expressed in mycobacteria using the expression vector p19Kpro as described previously [31]. To express WT and acetylation-defective mutant of PhoP, we cloned the entire *phoPR* operon with its 200-bp promoter region using ScaI and HindIII sites of promoter-less **pST-HiT** [44]. In this construct, we next introduced mutations in lysine residues (K195 and K197) of PhoP using two-stage overlap extension method [45]. The resulting wild-type and mutant constructs were verified by DNA sequencing, and expressed from a *phoPR*-KO background. The sequence of oligonucleotides used for amplification/cloning and the plasmids used for expression are listed in Table S1.

### Mycobacterial protein fragment complementation (M-PFC) assays

This assay is based on the principle that if two mycobacterial proteins fused to two different domains of dihydrofolate reductase (DHFR) enzyme interact, the functional reconstitution of the enzyme would allow *M. smegmatis* cells (expressing the relevant protein pair) to grow in the presence of Trimethoprim, a substrate of DHFR. *M. tuberculosis phoP* and 11 acetyltransferases were PCR amplified, cloned in pUAB400 (Kan^r^) and pUAB300 (Hyg^r^), respectively and expressed in *M. smegmatis* mc^2^155. The primer pairs used for cloning related to M-PFC assays and the relevant plasmid constructs used in this study are listed in Table S2. Growth of the co-transformants was monitored for few days on 7H11/Kan/Hyg plates supplemented with 10 μg/ml of Trimethoprim (TRIM). While growth on TRIM plates suggests interactions between two co-expressing proteins, empty vectors were included as negative controls and a pair of known interacting partner PhoP/PhoR proteins served as a positive control. All of the bacterial strains grew well in the absence of TRIM.

### Expression and purification of proteins

*M. tuberculosis* PhoP, DosR, and acetyl transferases were over-expressed in *E. coli* BL21(DE3) as 6xHis-tag proteins, and purified by Ni-affinity chromatography as described previously [42, 43]. In all cases, the purity of proteins was checked by SDS-polyacrylamide gel electrophoresis, and protein concentrations were determined by Bradford reagent using BSA as a calibration standard.

### *In vitro* acetylation assay

*In vitro* acetylation reactions were carried out in a buffer comprising 50 mM Tris-HCl (pH 8.0), 100 mM NaCl and 10 mM sodium butyrate, followed by the addition of 1 µg acetyltransferases, 2 µg PhoP, 0.5 mM acetyl CoA and 1mM cAMP in a reaction volume of 20 μl. The reaction mix was incubated at 25°C for 2 h, followed by addition of loading dye containing SDS to stop the reaction, and the products were resolved by SDS-PAGE. Acetylation was assessed by Western blot analysis using anti-acetyl antibody.

### Mass spectroscopy

Purified PhoP protein was loaded in 10% SDS-PAGE gel and stained using Coomassie blue G250, protein bands were excised and standard protocol was followed for preparing samples for mass spectrometry. Prior to tryptic digestion of the protein overnight using mass-spec grade Trypsin (Promega), protein was reduced and alkylated using dithiothreitol and iodoacetamide, respectively. Next, the resulting tryptic peptides were treated with 60% acetonitrile and 0.1% TFA. For LC-MS acquisition, peptide clean-up was performed using Zip-tip according to manufacturer’s recommendation. A Triple TOF 600 mass spectrometer (AB SCIEX, USA) coupled to nano-LC system was used to acquire samples and the results were analysed using Protein pilot software 5.0.1 (SCIEX, USA) using the Paragon algorithm. Analyst TF 1.7.1 software was used for data acquisition using optimized source parameters. During LC-MS/MS acquisition a biological modification with acetylation emphasis was enabled in ID focus. The detected protein threshold was set at >0.05 (10.0%), false discovery rate (FDR) analysis was enabled, only proteins identified with a global FDR of 1% were considered, and peptides with lysine residues modified due to acetylation were identified.

### RNA isolation

Total RNA from *M. tuberculosis* cultures grown with or without specific stress were isolated as described previously [41]. Briefly, 25 ml of bacterial culture was grown to mid-log phase (OD_600_ ∼ 0.4 to 0.6) and combined with 40 ml of 5 M guanidinium thiocyanate (GTC) solution containing 1% β-mercaptoethanol and 0.5% Tween 80. Cells were pelleted by centrifugation, and lysed by re-suspending in 1 ml Trizol (Ambion) in the presence of Lysing Matrix B (100 µm silica beads; MP Bio) using a FastPrep-24 bead beater (MP Bio) at a speed setting of 6.0 for 30 seconds. The procedure was repeated for 2-3 cycles with intermittent cooling on ice in between pulses. Cell lysates were centrifuged at 13000 rpm for 10 minutes, supernatant collected and RNA isolation carried out using Direct-Zol^TM^ RNA isolation kit (ZYMO). To remove genomic DNA, following extraction RNA was treated with RNase -free DNase I (Promega) for 30 minutes, and integrity was assessed by measuring A_260_ and A_280_ in a spectrophotometer. RNA samples were further verified for intactness of 23S and 16S rRNA using formaldehyde-agarose gel electrophoresis, and Qubit fluorometer (Invitrogen). In all cases RNA preparations were free from genomic DNA.

### Quantitative Real-Time PCR

In this assay, total RNA obtained from *M. tuberculosis* were used to perform RT-qPCR using Superscript III platinum-SYBR green one-step kit (Invitrogen). cDNA was generated from 50 ng of RNA using Superscript III platinum enzyme, SYBR mix and primers were added together and amplified in an Applied Biosystems real-time PCR detection system. Sequences of oligonucleotide primers used in RT-qPCR and ChIP-qPCR experiments are listed in Table S3. A standard curve was generated for each pair of primers using serially diluted RNA samples to determine the PCR efficiency, and in all cases PCR efficiency was within 95-105%. Use of platinum Taq DNA polymerase (Invitrogen) as control reactions ensured absence of genomic DNA in our RNA preparations, and endogenously expressed *M. tuberculosis 16S rDNA* served as the internal control. Fold difference in gene expression was determined using ΔΔC_T_ method [46].

### ChIP-qPCR

ChIP experiments were performed using anti-PhoP antibody as detailed previously [41]. Briefly, mycobacterial cells were grown to mid exponential phase (OD_600_ ≈0.4-0.6) and formaldehyde was added to a final concentration of 1%. After incubation of 20 minutes, glycine was added to a final concentration of 0.5 M to quench the reaction and incubated for further 10 minutes. Cross-linked cells were harvested by centrifugation, and washed in immunoprecipitation (IP) buffer [50 mM Tris (pH 7.5), 150 mM NaCl, 1 mM EDTA, 1% Triton X-100, 1 mM PMSF and 5% glycerol]. Further details of processing of the IP DNA, shearing to an average size of ∼ 500-bp, and final recovery after ethanol precipitation has been described previously [41]. Enrichment due to recruitment of the regulator *in vivo* was determined using appropriate dilutions of IP DNA in a reaction buffer containing SYBR green mix (Invitrogen), 2 µM PAGE-purified primers and one unit of Platinum Taq DNA polymerase (Invitrogen). qPCR signal from a mock control lacking antibody was measured to determine the efficiency of recruitment. In all cases, melting curve analysis confirmed amplification of a single product. Specificity of PCR-enrichment from the identical IP samples was verified using *16S rDNA/rpoB*-specific primers. Each data was recorded in duplicate qPCR measurements using at least two bacterial cultures.

### Isolation of *ex vivo* peritoneal macrophages and infection experiments

Experiments pertaining to mice macrophages were in accordance with Institutional regulations after review of protocols and approval by the Institutional Animal Ethics Committee (IAEC/21/03). Mice were maintained and bred in the animal house facility of CSIR-IMTECH. 4 C57BL/6 mice (6-8 weeks) were used for isolation of peritoneal macrophages. Briefly, 4% thioglycollate solution was injected into the peritoneum cavity of mice. On attaining 4th day, elicited peritoneal macrophages were isolated from the peritoneum cavity using serum-free media and plated using complete RPMI 1640 media for adherence. Prior to experiment, the nonadherent cells were removed by washing and the adherent macrophages were used for infection experiments as described previously [47, 48]. The macrophages were infected with *M. tuberculosis* strains at an MOI of 1:10, the cells were lysed at appropriate time points, and CFU was enumerated.

### Macrophage Infections

RAW264.7 macrophages were infected with titrated cultures of WT-H37Rv, and mutant mycobacterial strains as described previously [31]. While phenolic auramine solution was used to stain *M. tuberculosis* H37Rv strains, the cells were stained with 150 nM Lyso-Tracker Red DND 99 (Invitrogen). Cells were fixed, analysed using confocal microscope (Nikon, A1R), and processing of digital images were carried out with IMARIS imaging software (version 9.20). Details of the experimental methods and the laser/detector settings were optimized using macrophage cells infected with WT-H37Rv as described [2]. A standard set of intensity threshold was made applicable for all images, and percent bacterial co-localization was determined by analyses of at least 50 infected cells originating from 10 different fields of each of the three independent biological repeats.

## Acknowledgements

We are grateful to G. Marcela Rodriguez and Issar Smith (PHRI, New Jersey Medical School - UMDNJ) for *phoP*-KO, and the complemented mutant strains, and Adrie Steyn (University of Alabama) for pUAB300/pUAB400 plasmids. We thank members of the Institutional animal facility (iCARE) for their help with approval of our project from the Institutional Animal Ethics Committee. This study received financial support from intramural grants of CSIR-IMTECH (OLP-0170), and CSIR (MLP-0049). P.P., K.M., R.U., B.B. and H. G., were supported by CSIR pre-doctoral fellowships. B.T., received financial support from CSIR.

