## Supplemental Information for "Context-dependent acetylation of the virulence regulator PhoP accounts for carbon-source specific intracellular growth program of *Mycobacterium tuberculosis*"

Context-dependent acetylation of the virulence regulator PhoP accounts for carbon-source specific mycobacterial growth inhibition under acidic conditions

Partha Paul^1^, Khushboo Mehta^1,4^, Rajat Ujjainiya^2^, Bhanwar Bamniya^1^,

Bhuwaneshwar Thakur^1^, Harsh Goar^1,3,^ Dibyendu Sarkar^1,4,*^

^1^ CSIR-Institute of Microbial Technology, Sector 39 A, Chandigarh 160036, India

^2^ CSIR-Institute of Genomics and Integrative Biology, New Delhi 110025, India

^3^ Present address: Department of Medicine, Division of Hematology-Oncology,

UT Southwestern Medical Center, Dallas, TX 75235

^4^ Academy of Scientific and Innovative Research (AcSIR), Ghaziabad, India

Running title: Mycobacterial growth inhibition and PhoP acetylation

*Address correspondence to: Dibyendu Sarkar, CSIR-Institute of Microbial Technology

Key Words: Acidic pH; Carbon source utilization; Mycobacterial growth inhibition;

PhoP acetylation; Virulence regulator PhoP

**Table S1**

Sequences of oligonucleotide primers and plasmids used for cloning reported in this study

| Primers^a^ | Sequence or description (5' - 3') | Reference |
| --- | --- | --- |
| FPRv0133 | AATAATCATATGATGACCCCGCA | This study |
| RPRv0133 | AATAATAAGCTTCTACCGAGGCTC | This study |
| FPRv0998 | AATAATCCATGGTTGGACGGGATAGCC | This study |
| RPRv0998 | AATAATAAGCTTTCAGCCGACGGCCTCGAT | This study |
| FPRv2851 | AATAATGGATCCATGACCGAA | This study |
| RPRv2851 | AATAATAAGCTTTCACGGCCGCT | This study |
| FPRv3027 | AATAATGGATCCATGAGCATCGCTT | This study |
| RPRv3027 | AATAATAAGCTTTCATCGCGCGT | This study |
| FPdosRstart | AATAATCATATGATGGGAAGCGCCGA | [1] |
| RPdosRstop | AATAATGGATCCTCATCGAGCACCCA | [1] |
| FPphoP19K | GTTTGCGGATCCATGCGGAAAGGGGTTGAT | [2] |
| RPphoP19K | GGTGGTCTGCAGTCAGTGGTGGTGGTGGTGGTGTCGAGGCTCCCGCAG | [2] |
| FPphoP promoter | AATAATAGTACTGCGAGCCCCG | This study |
| RPphoR | AATAATAAGCTTTCAGGGCGGCC | This study |
| FPphoPK195R | CCGTGCTGAGCAGGCCTAAGATTC | This study |
| RPphoPK195R | GAATCTTAGGCCTGCTCAGCACGG | This study |
| FPphoPK197R | GAGCAAGCCTAGGATTCTCGACCAC | This study |
| RPphoPK197R | GTGGTCGAGAATCCTAGGCTTGCTC | This study |
| Plasmids |  |  |
| pET-28a^a^ | *E. coli* cloning vector | Novagen |
| pET-*rv2851* | His_6_-tagged Rv2851expression plasmid | This study |
| pET-*rv3027* | His_6_-tagged Rv3027 expression plasmid | This study |
| pET-28b^a^ | *E. coli* cloning vector, Kan | Novagen |
| pET-*rv0133* | His_6_-tagged Rv0133 expression plasmid | This study |
| pET-*rv0998* | His_6_-tagged Rv0998 expression plasmid | This study |
| pET-*dosR* | DosR residues 1–217 cloned in pET28b | [1] |
| p19Kpro^b^ | Mycobacterial expression vector | [3] |
| p19Kpro-*phoP*His | PhoP residues 1–247 cloned in p19Kpro with His_6_-tag | [2] |
| pST-HiT^b^ | Mycobacterial integrative expression vector | [4] |
| pST-PhoPR | PhoPR operon comprising 200bp promoter cloned in pST-HiT | This study |
| pST-*phoP*K195R*phoR* | Lys195 mutated to Arg in *phoP* of pST-*phoPR* | This study |
| pST-*phoP*K197R*phoR* | Lys197 mutated to Arg in *phoP* of pST-*phoPR* | This study |

^a^kanamycin resistance; ^b^ hygromycin resistance

FP, forward primer; RP, reverse primer

**Table S2**

Sequences of oligonucleotide primers and plasmids used in M-PFC experiments

| ^a^Primers | Sequence or description (5’-3’) | Reference |
| --- | --- | --- |
| FPphoP | AATAATCAATTGATGCGGAAAGGG | [5] |
| RPphoP | AATAATAAGCTTTCATCGAGGCTC | [5] |
| FPphoR | AATAAACTGCAGATGGCCAGACAC | [5] |
| RPphoR | AATAATAAGCTTTCAGGGCGGCC | [5] |
| FPRv0133 | AATAATCTGCAGATGACCCCGCA | This study |
| RPRv0133 | AATAATAAGCTTCTACCGAGGCTC | This study |
| FPRv0428 | AATAATGGATCCATGGTCTCGTG | This study |
| RPRv0428 | AATAATAAGCTTCTAGAAGGTATC | This study |
| FPRv0730 | AATAATCTGCAGATGCATGGCGCA | This study |
| RPRv0730 | AATAATAAGCTTCTACCGGCCGGCA | This study |
| FPRv0998 | AATAATCCATGGTTGGACGGGATAGCC | This study |
| RPRv0998 | AATAATAAGCTTTCAGCCGACGGCCTCGAT | This study |
| FPRv1347c | AATAATGGATCCATGACCAAACCC | This study |
| RPRv1347c | AATAATAAGCTTTTACGCAGCCGT | This study |
| FPRv2170 | AATAATGGATCCTTGGCGATATTC | This study |
| RPRv2170 | AATAATAAGCTT TTAGAGCGGTAG | This study |
| FPRv2416c | TATATAGGATCCGTGACTGTGACCCTG | This study |
| RPRv2416c | CTAGCACTGCAGTCAGAACTCGAACGCG | This study |
| FPRv2669 | AATAATGGATCCGTGACCGAC | This study |
| RPRv2669 | AATAATAAGCTTTCATACAAG | This study |
| FPRv2775 | AATAATGGATCCATGCACTATC | This study |
| RPRv2775 | AATAATAAGCTTTTACCCGGGCGG | This study |
| FPRv2851 | AATAATGGATCCATGACCGAA | This study |
| RPRv2851 | AATAATAAGCTTTCACGGCCGCT | This study |
| FPRv3027 | AATAATGGATCCATGAGCATCGCTT | This study |
| RPRv3027 | AATAATAAGCTTTCATCGCGCGT | This study |
| Plasmids | Descriptions | References |
| pUAB400^a^ | Integrative expression plasmid, Kan^r^ | [6] |
| pUAB-*phoP* | pUAB400 expressing PhoP (aa 1-247) | [7] |
| pUAB300^b^ | Episomal expression plasmid, Hyg^r^ | [6] |
| pUAB300-*phoR* | pUAB300 expressing PhoR (aa 1-485) | [7] |
| pUAB300-*rv0133* | pUAB300 expressing Rv0133 (aa 1-579) | This study |
| pUAB300-*rv0428* | pUAB300 expressing Rv0428 (aa 1-579) | This study |
| pUAB300-*rv0730* | pUAB300 expressing Rv0730 (aa 1-574) | This study |
| pUAB300-*rv0998* | pUAB300 expressing Rv0998 (aa 1-861) | This study |
| pUAB300-*rv1347c* | pUAB300 expressing Rv1347c (aa 1-505) | This study |
| pUAB300-*rv2170* | pUAB300 expressing Rv2170 (aa 1-568) | This study |
| pUAB300-*rv2416c* | pUAB300 expressing Rv2416c (aa 1-502) | This study |
| pUAB300-*rv2669* | pUAB300 expressing Rv2669 (aa 1-447) | This study |
| pUAB300-*rv2775* | pUAB300 expressing Rv2775 (aa 1-411) | This study |
| pUAB300-*rv2851* | pUAB300 expressing Rv2851 (aa 1-476) | This study |
| pUAB300-*rv3027* | pUAB300 expressing Rv3027 (aa 1-510) | This study |

FP, forward primer; RP, reverse primer

**Table S3**

Sequences of oligonucleotide primers used in RT-PCR and ChIP experiments reported in this study

| Primers | Sequence (5’ to 3’) | Reference |
| --- | --- | --- |
| FPaprART | TTGACCATGACAGCGAGTGT | [5] |
| RPaprART | TTGGACAGAAATGCAGGATG | [2] |
| FPlipFRT | TAGTGGCCATCTCTCCGTTG | This study |
| RPlipFRT | AGCGGCTCATAGAGGTCTTC | This study |
| FPmsl3RT | GTGAAAACAAACTTCGGTCAC | [8] |
| RPmsl3RT | ACAAAGAGTTCAGTGTCAATCTCAG | [8] |
| FPpks2RT | GTTGTGGAAGGCGTTGTTAC | [8] |
| RPpks2RT | GTCGTAGAACTCGTCGCAAT | [8] |
| FPwhiB3RT | TGGACTCATCGATGTTCTTCC | [5] |
| RPwhiB3RT | TAGGGCTCACCGACCTCTAA | [5] |
| FPrpoBRT | GGAGGCGATCACACCGCAGACGTT | [9] |
| RPrpoBRT | CCTCCAGCCCGGCACGCTCACGT | [9] |
| FP16SrDNART | CTGAGATACGGCCCAGACTC | [9] |
| RP16SrDNART | CGTCGATGGTGAAAGAGGTT | [9] |
| FPaprAup | CAGGTTTGCCTCCCAGCCGC | This study |
| RPaprAup | GGGCCGCCGGCTTTGTCAC | This study |
| FPlipFup | AAACTCCCGCATAGAACACCT | This study |
| RPlipFup | GCTCCAACCAGACGCAGCC | This study |
| FPpks2up | AATAATGGATCCGAAGCGTCAGACTACCGG | [8] |
| RPpks2up | AATAATGGTACCTATCTGCACCAGTGCCTG | [4] |
| FPwhib3up | GAATGCGGCGCAGATGAACG | [5] |
| RPwhib3up | GTAGCTGCTCCGGCTGTGG | [5] |
| FPespAup | CGTGATCTTGATACGGCTCG | [2] |
| RPespAup | GTTGTTGGTACCCTCGGCAAGATCGGC | [2] |
| FP16SrDNAup | CTGAGATACGGCCCAGACTC | [7] |
| RP16SrDNAup | CGTCGATGGTGAAAGAGGTT | [7] |

FP, forward primer; RP, reverse primer

**Figure legends**

**Fig. S1: *phoP* is required for utilization of acetate as a carbon source under normal conditions (pH 7.0)**. (A-B) In this experiment, indicated mycobacterial strains were grown in a minimal media containing either glycerol or acetate, under normal conditions (pH 7.0). Mycobacterial strains were inoculated at an initial OD_600_ of 0.05 at specific media, as described in the Methods, and growth was monitored at different time points. To enumerate CFU values, cells were harvested both at the onset of growth (0 day) and on the 6^th^ day. The values represent average of two biological repeats with two technical repeats (*P<0.05; ***P<0.001).

**Fig. S2: Screening to probe PhoP interacting mycobacterial acetyl transferases.** (A-G) M-PFC experiments co-expressing *M*. *tuberculosis* PhoP and acetyl transferases were used as a screen using *M*. *smegmatis* as the surrogate host. Co-expression of pUAB400-*phoP*/pUAB300-acetyl transferases (indicated) pair (columns 1 and 5) relative to empty vector controls, pUAB400-*phoP*/pUAB300 (columns 2 and 6), or pUAB400/pUAB300-acetyl transferases (columns 3 and 7), supporting *M*. *smegmatis* growth in the presence of TRIM is suggestive of specific interaction. In each case the three spots moving downwards in each panel represent spotting of cells at three different dilutions, namely undiluted, 10-fold and 100-fold dilutions, respectively. While co-expression of pUAB400-*phoP*/pAUB300-*phoR* (columns 4 and 8) in *M*. *smegmatis* displaying growth in the presence of TRIM served as a positive control, growth of all the strains in absence of TRIM (columns 5–8) validated the assay.

**Fig. S3: Sequence alignment of PhoP and members of the family of proteins.** (A) Multiple sequence alignment of full-length PhoP from various bacterial species highlights the conserved lysine residues (indicated in blue colour). (B-C) To express *phoPR* ORF, the entire operon along with 200-bp promoter was cloned in promoter-less pST-HiT [4]. The construct was also engineered further to introduce mutations in K195 and K197 of PhoP sequence using two-stage overlap extension method. The resulting wild-type and mutant constructs were expressed in a *phoPR*-KO mutant background. Mycobacteria harbouring wild-type or indicated *phoP* mutants were inoculated at an initial OD_600_ of 0.05 in the presence of acetate as a carbon source, and growth curves were compared under (B) acidic and (C) normal conditions.

Figure S1

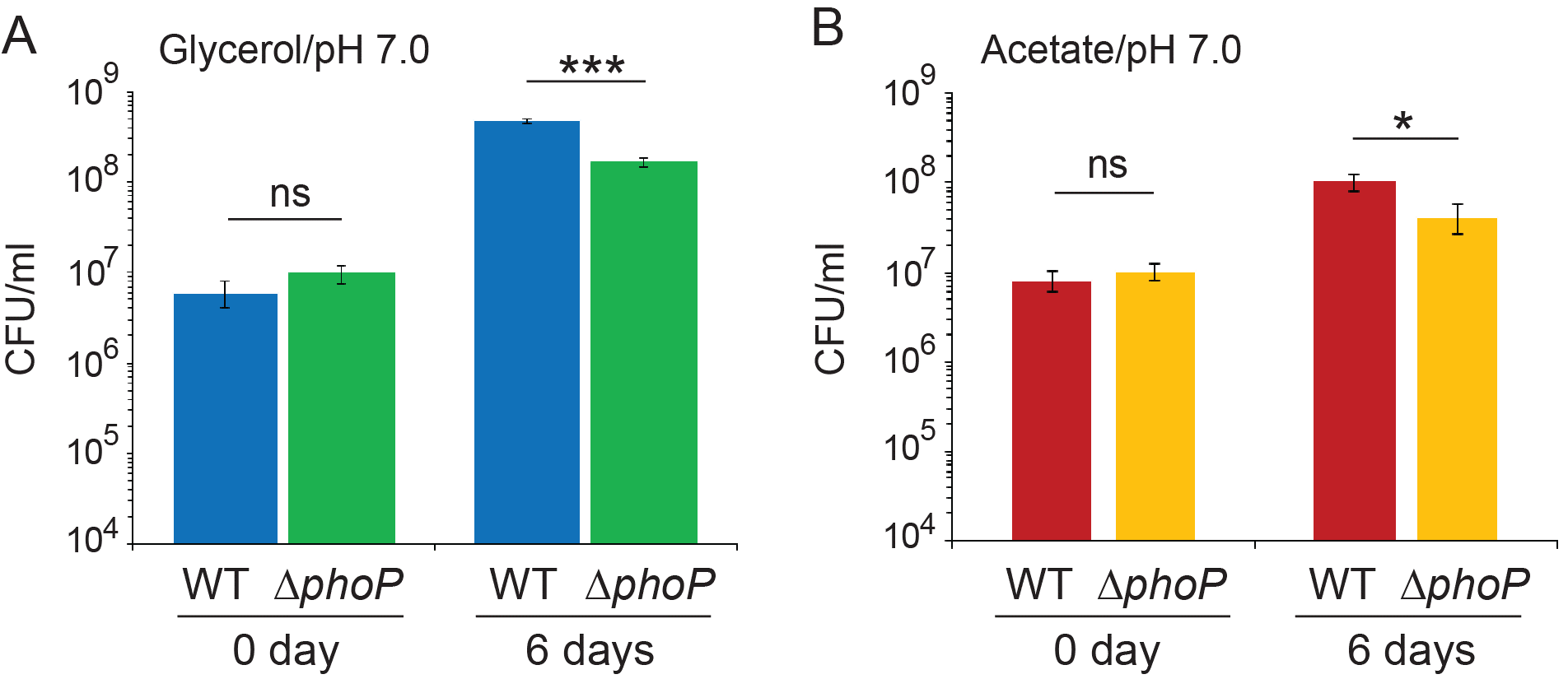

Figure S2

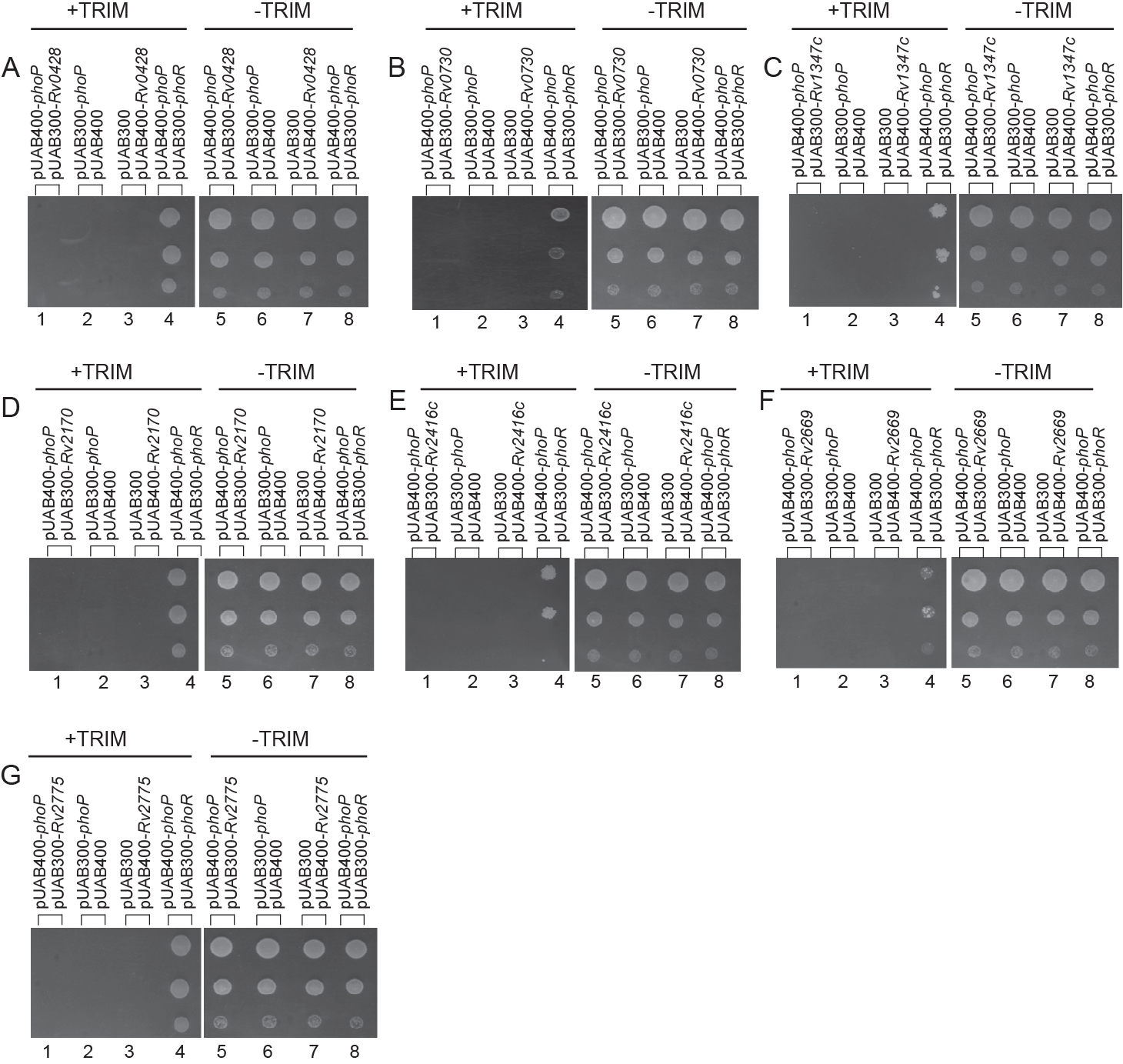

Figure S3

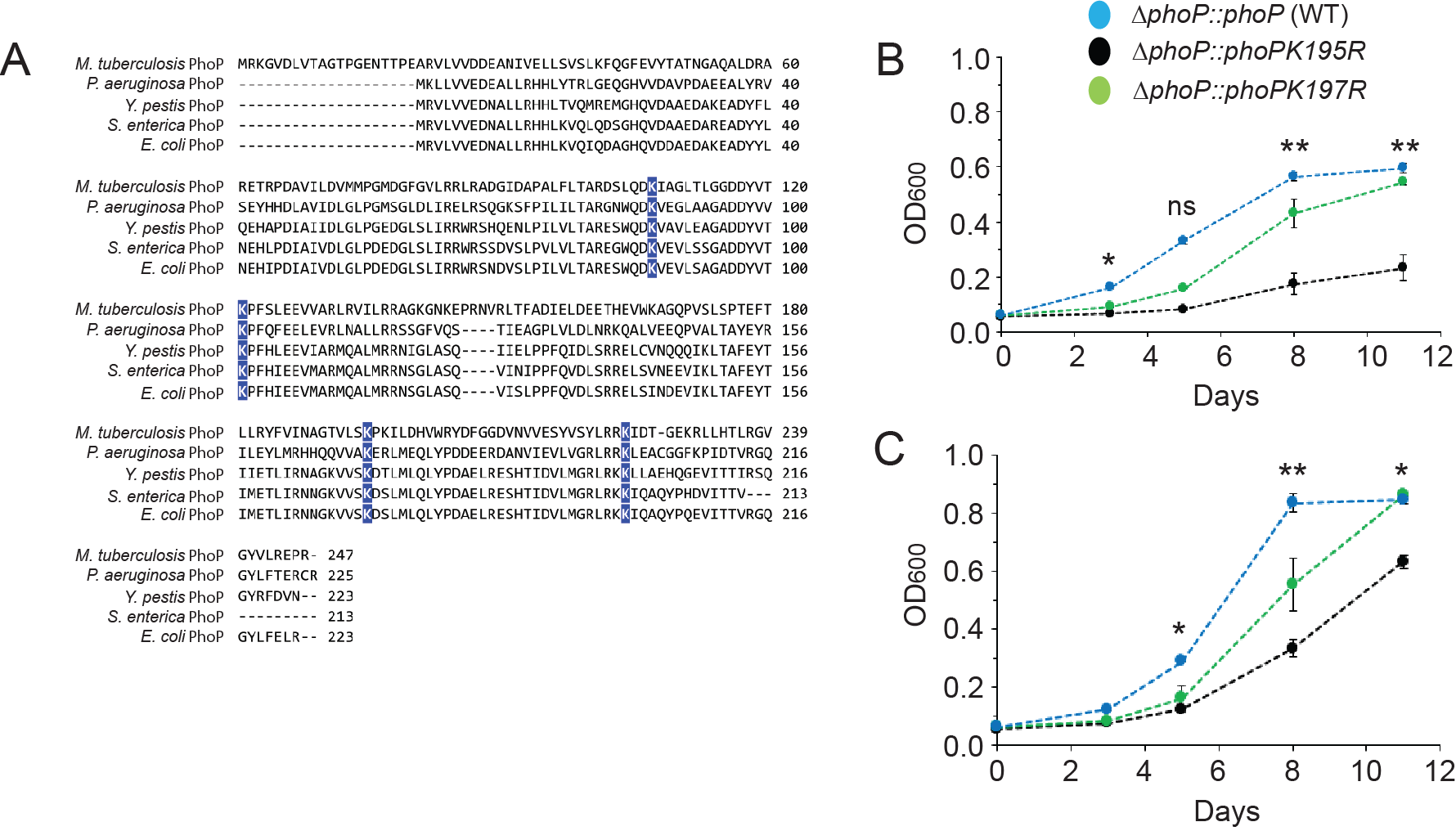
